# Microsatellite Analysis of Genetic Differentiation Among Populations of the Malaria Vector, *Anopheles funestus*, across Mainland Tanzania Reveals Contrasted Patterns of Geographic Isolation and Gene Flow

**DOI:** 10.64898/2025.12.14.694266

**Authors:** Salum Abdallah Mapua, Badara Samb, Ismail Hassan Nambunga, Prisca Asiimwe Kweyamba, Gustav Mkandawile, Hamis Bwanaly, Emmanuel Wilson Kaindoa, Joel Ouma Odero, John Paliga Masalu, Najat Feruz Kahamba, Emmanuel Elirehema Hape, Roberto Galizi, Nicodem James Govella, Fredros Oketch Okumu, Frederic Tripet

## Abstract

**Background:** Despite Anopheles funestus s.s. being a highly competent and widespread malaria vector in Africa, its population structure remains largely understudied in many countries, including Tanzania. Herein, we examine the genetic diversity, geographic isolation, and gene flow of An. funestus populations across 10 administrative regions in mainland Tanzania.

**Methods:** We employed 12 previously used microsatellite DNA markers to describe genetic diversity, isolation by distance, and gene flow patterns among 10 An. funestus s.s. populations (n = 654) and one An. parensis population (n = 28), used as an outgroup, sampled across 10 regions in mainland Tanzania.

**Results:** Overall, allelic richness (*N*a) and genetic diversity (*H*_S_) did not differ significantly among populations. Although some loci and populations showed significant departures from Hardy-Weinberg equilibrium, the patterns were not indicative of sub-structuring within locations. Pairwise genetic divergence (*F*_ST_) values indicated clear separation between *An. parensis* and *An. funestus* s.s., with values exceeding 0.2, consistent with species-level differentiation. Among *An. funestus* s.s. populations, the highest divergence was observed between southeastern coastal populations (Mtwara, Ruvuma, Lindi) and inland populations, with *F*_ST_ values up to 0.288. There was no evidence of isolation by distance. Instead, patterns of genetic divergence suggested connectivity across the Rift Valley and heterogeneity among southeastern populations. Neighbour-joining analysis and Bayesian genotype clustering identified three distinct population groups: (i) *An. parensis* (Dodoma), (ii) a genetically distinct *An. funestus* s.s. population from Mtwara, and (iii) a more homogeneous cluster comprising the remaining *An. funestus* s.s. populations. Notably, the Mtwara population appeared highly differentiated, with divergence approaching that between *An. Funestus* s.s. and *An. parensis*, supporting its distinctiveness but not undermining the role of *An. parensis* as an outgroup.

**Conclusion:** Except for the Mtwara population, whose status will need to be clarified through whole genome sequencing, moderate genetic divergence was found among *An. funestus* s.s. populations across Tanzania, despite geographical separation and the Rift Valley. The observed genetic structure suggests that anthropogenic gene flow may play a key role in shaping population divergence. Future studies should aim to delineate the effects of local adaptation from recent gene flow to further explore these dynamics.

## Background

*Anopheles gambiae* sensu stricto, *An. arabiensis*, and *An. funestus* s.s. are the primary malaria vectors in Eastern Africa, contributing to 249 million cases and 608,000 deaths annually in sub-Saharan Africa [1]. These species belong to the *An. gambiae* complex and *An. funestus* group, with the *An. funestus* group comprising at least 13 sibling species, including *An. funestus* s.s., *An. parensis*, *An. rivulorum*, and others [2–9]. *An. funestus* s.s. is the most competent vector in the group [10] while species like *An. rivulorum*, *An. leesoni*, and *An. parensis* play a lesser role in *Plasmodium falciparum* transmission [3,5–7,9].

In Eastern and Southern Africa, *An. funestus* has become an increasingly dominant malaria vector over the past two decades [10], contributing to 90% of malaria cases in south-eastern Tanzania [11,12]. Its growing impact is linked to high insecticide resistance [13], reducing the efficacy of ITNs and IRS, as well as its strong preference for human hosts [14]. Despite its importance, the ecology and population structure of *An. funestus* s.s. remain poorly studied in many parts of Africa. In Tanzania, genetic studies on its behavior and resistance spread are limited, despite recent discoveries of voltage-gated sodium channel *(Vgsc*) gene mutations conferring knockdown resistance [15]. With emerging genetic control strategies, such as genetically modified mosquito releases [16], understanding *An. funestus* population structure is crucial for optimizing vector control interventions.

In the past, molecular techniques such as analysis of chromosomal inversions, mitochondrial DNA, restriction fragment length polymorphisms, and the commonly used microsatellite DNA markers have been employed to study population structure of *An. funestus* mosquitoes [17]. Although recent genomic approaches such as single nucleotide polymorphism (SNP) genotyping and whole-genome sequencing (WGS) offer unprecedented resolution, microsatellites remain valuable for studying recent gene flow and fine-scale population structure due to their high allelic diversity and mutation rates [18–21].

In this study, we examine the population genetic diversity, geographic isolation, and patterns of gene flow in *An. funestus* s.s. populations across 10 administrative regions covering most of mainland Tanzania. Clustering and Bayesian genotype assignments analyses were used to identify outlying populations in terms of genetic divergence, which was further facilitated by comparison with an *An. parensis* population used as outgroup. The results constitute an important preliminary study of *An. funestus* s.s. describing population interconnectivity and discontinuities across Tanzania with implication for the spread of insecticide resistance loci and vector control tools relying on the spread of genes such as GM mosquito release strategies.

## Methods

### Study area

*An. funestus* mosquitoes were collected from 10 administrative regions across Tanzania mainland on both sides of the eastern arm of the Great Rift Valley (Fig. 1, Additional file 1). The collection sites were chosen to cover varied climatic zones with temperature ranging from 20°C to 30°C and annual rainfall ranging from 550 to 3690 mm [22]. These were the hot and humid coastal plain (i.e. Pwani, Tanga, Lindi and Mtwara), the semi-arid central plateau (i.e., Dodoma, Kigoma, Katavi and Rukwa), the high rainfall lake regions (i.e., Kagera), and the cooler highlands (i.e., Morogoro and Ruvuma). Subsistence farmers constituted most of households in the villages that were sampled [23,24]. Malaria prevalence under the age of 5 ranges from less than 1% in Dodoma to 20% in Mtwara [25].

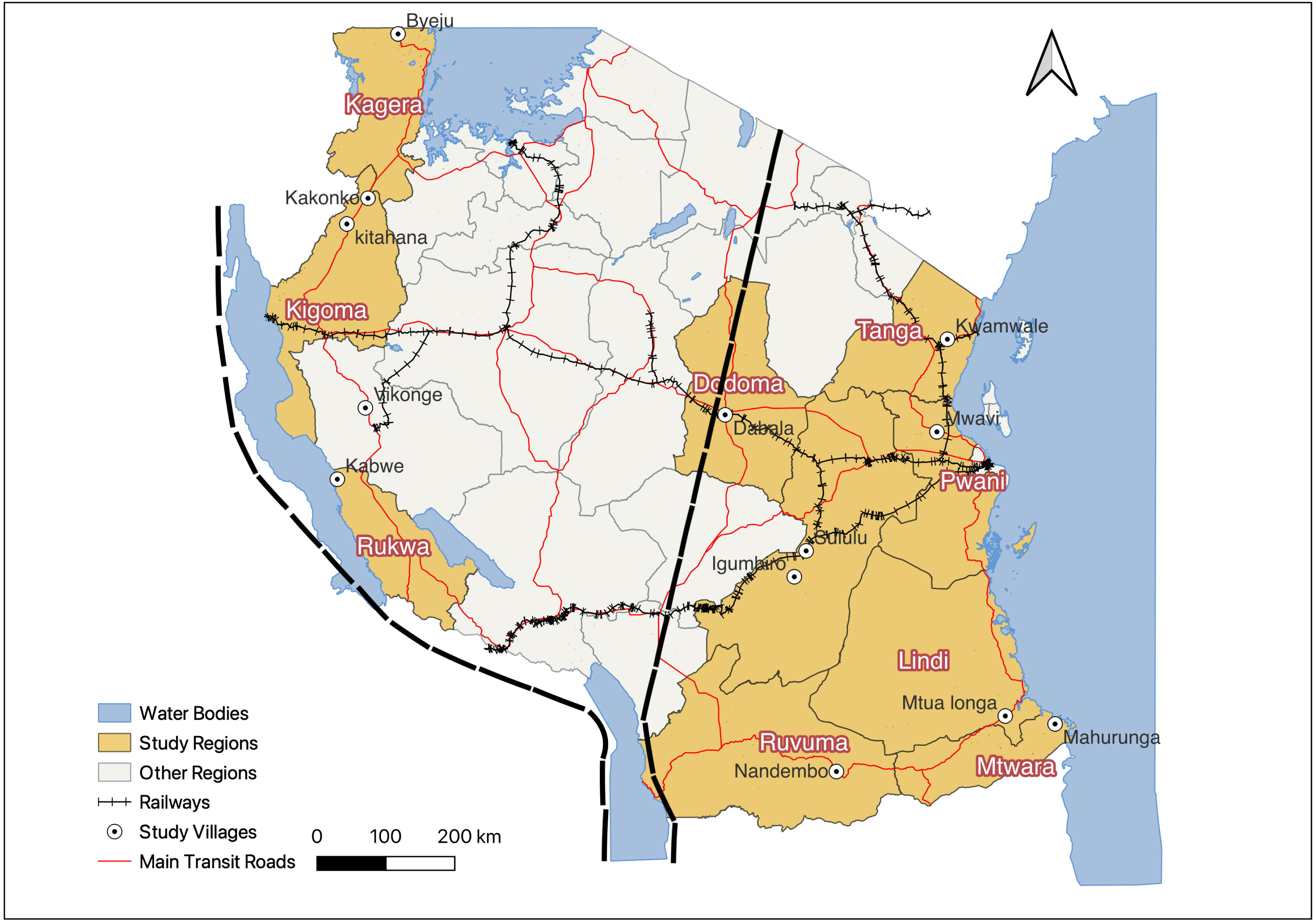

### Mosquito collection

Mosquito collections were conducted between December 2018 and December 2022, from at least two houses per district, upon consent from household heads, using CDC light traps for indoor host-seeking [26] and Prokopack aspirators for indoor-resting mosquitoes [27]. The mosquitoes were morphologically identified to species complex level and *An. funestus* group retained for molecular characterization using PCR. Female *An. funestus* were individually stored in Eppendorf tubes containing 80% ethanol. Additionally, in certain regions such as Dodoma, Tanga, and Morogoro, where adult *An. funestus* sampling was insufficient, larval collections were performed using standard larval dippers [28]. The larvae were reared to adults, sorted by species, and processed as described [29].

### Genomic DNA extraction

Genomic DNA was extracted using the DNAzol method [30]. Mosquito heads and thoraces were placed individually in a 96-well plate with 100 µl of DNAzol and three sterile beads. A negative control (DNAzol and beads only) was included. Samples were homogenized (Bead Ruptor 96, OMNI International) for 12 cycles of 30 seconds at 22 Hz, then centrifuged at 4000 rpm for 15 minutes. The supernatant was mixed with ethanol, washed twice with 75% ethanol, and dried before elution in 50 µl of TE buffer. DNA concentration was measured using a Nanodrop 1000 spectrophotometer (Thermo Fisher Scientific).

### Molecular identification of *Anopheles funestus* sibling species

The extracted genomic DNA was used in a cocktail of species-specific primers to identify the sibling species within the *An. funestus* group, following the method described by Koekemoer *et al*. [31] with a small modification to include a primer for *Anopheles rivulorum-like* [8,32]. Whilst other sibling species of the *An. funestus* group were identified from the study area, only *An. funestus* s.s. which was the dominant species in our collections was used for further genetic analyses. Additionally, *An. parensis* individuals from Dodoma, central Tanzania, were included to act as a genetically distinct outgroup in our analyses.

### Microsatellite genotyping

Eighteen microsatellite markers previously used in population genetic studies [21,33] were considered to describe the population structure of the *An. funestus* s.s. across mainland Tanzania (Fig. 1, Additional file 2). These markers were selected based on their spread across the entire *An. funestus* genome, polymorphism and for having no evidence of null alleles [21]. They were amplified from extracted genomic DNA in a 20 µl PCR mix. The reaction mix contained 10 µl of PCRBIO HS Taq Mix Red (PCR Biosystems, Aztec House, London, UK), 0.5 µl of each of the forward and reverse primers of the specific marker at a concentration of 0.3 µM, 8 µl of double distilled water and 1 µl of sample DNA. The forward primer of each marker was labelled in 5’ prime end with either by FAM (i.e., Fluorescein amidite), Yakima Yellow or ATTO 565 fluorescent markers to allow multiplex electrophoresis (Eurofins Dye set EF-01). A PCRmax thermal cycler (PCRmax Limited, ST, UK) was used with the following thermal cycling conditions for 40 cycles: denaturation at 95°C, annealing varying between 51 and 61.5°C depending on the microsatellite marker, and extension at 72°C. Three microliters of the resulting PCR products were electrophoresed on a 1.5% agarose gel stained with Gel Red, running for 120 min at 80 volts. The amplified genomic DNA fragments were visualized using a gel documentation system (GBOX-Chemi-XRQ, Syngene). Additionally, amplified products were diluted if necessary and pooled before being shipped to Eurofins Genomics (Ebersberg, Germany) for genotyping on an AB3130 sequencer (Applied Biosystems, Foster, California, USA). Alleles were scored and sized relative to an internal size standard using GENEMAPPER version 5.0 [34].

### Selection of best microsatellite loci

For each locus, allele sizes generated by GENEMAPPER were double-checked manually to ensure compatibility with their tandem motif molecular sequence and with the binning of allele size used in previous studies [21]. The number of alleles (*N*a, allelic richness) expected heterozygosity (*H*_S_, genotypic diversity) and observed heterozygosity (*H*_O_) per loci and overall were computed using software FSTAT version 2.9.3 [35] and GENEPOP version 4.7.5 [36]. Population deviations from Hardy-Weinberg Equilibrium (HWE) per loci and overall were computed using GENEPOP version 4.7.5 [36] with Bonferroni adjusted *p*-values. To eliminate loci that did not conform to neutrality, microsatellite loci that significantly deviated from Hardy-Weinberg equilibrium in more than four out of the 11 populations were excluded from further analyses resulting in a set of 12 preferred microsatellite loci (Fig. 2, Additional file 3). Pairwise genotypic linkage disequilibrium (LD) between all loci was tested using the Expectation-Maximization algorithm in GENEPOP v4.7.5. No significant LD was detected after Bonferroni correction (adjusted α = 0.00076), confirming that the retained loci are independently segregating.

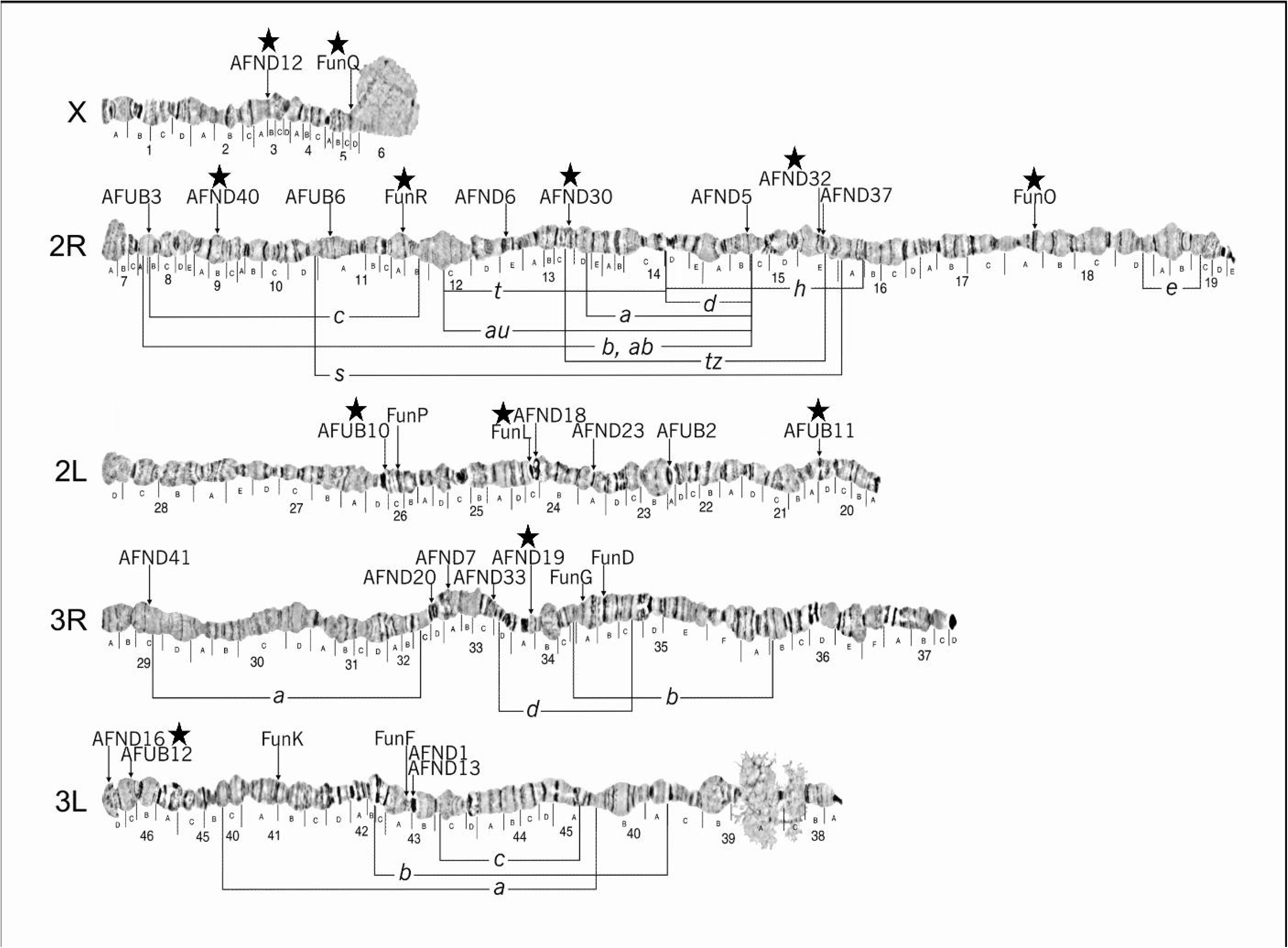

### Population genetics analysis

Estimates of genetic divergence among populations (*F*_ST_) and inbreeding coefficients (*F*_IS_) were estimated using GENEPOP version 4.7.5 [36] and FSTAT version 2.9.3 [35] respectively. Statistical significance of pairwise *F*_ST_ was assessed using Arlequin version 3.5 [37]. Pairwise *F*_ST_ estimates were used in the software MEGA 5 [38] to generate neighbour-joining tree, with the Dodoma population comprising *An. parensis*, a sibling species within the *An. funestus* group, included as an outgroup to root the analysis and highlight divergence from *An. funestus* s.s. populations. Isolation-by-distance between *An. funestus* s.s. populations was examined by testing the correlation between linearised *F*_ST_ (*F*_ST_ /(1- *F*_ST_) and log transformed geographical distances among subpopulations by Mantel test in GENEPOP version 4.7.5 [36] and by plotting them using a linear fit in the statistical software JMP^®^ version 17.2.0 (SAS Institute Inc., Cary, NC, 1989-2019). The Bayesian model-based clustering algorithm implemented in the software STRUCTURE version 2.2 was used to assign individual genotypes into inferred populations [39]. For each K value, we performed 10 independent replicate runs to assess consistency across runs. Every run consisted of 250,000 iterations following a burn-in period of 50,000 iterations. The model’s estimated likelihood values [Pr(X|K)], together with STRUCTURE bar plots and neighbor-joining trees, were used to identify the value of K most relevant to our dataset.

## Results

### Genetic diversity and heterozygosity among populations

A total of 654 *An. funestus* s.s. and 28 *An. parensis* individuals from 11 populations were genotyped. Among the 10 *An. funestus* s.s. populations, mean allelic richness (*N*a) varied from 4.7 in Lindi to 8.3 in Mtwara, with a mean *N*a across all populations of 6.6. In the *An. parensis* population from Dodoma *N*a was 7.5. There was however no statistically significant difference in mean *N*a between the 10 *An. funestus* s.s. populations (Kruskal-Wallis: χ^2^ = 14, df = 9, *p* = 0.122) (Additional file 4).

In *An. funestus* s.s., the mean expected heterozygosity (*H*_S_) across all loci per population ranged from 0.467 in Lindi to 0.703 in Pwani, while observed heterozygosity (*H*_O_) ranged from 0.453 in Rukwa to 0.781 in Pwani. The populations with next highest *H*_S_ were those of Kigoma (0.675) and Kagera. No significant differences in *H*_S_ or *H*_O_ were found across the 10 populations (Kruskal-Wallis: χ² > 16.3, df = 9, *p* < 0.050 in both cases). The mean *H*_S_ and *H*_O_ for *An. funestus* s.s. populations were 0.586 and 0.585, respectively; compared to 0.640 and 0.658 for *An. parensis*.

### Hardy-Weinberg equilibrium

A multi-locus analysis of Hardy-Weinberg equilibrium revealed significant heterozygote deficits in 9 out of 11 populations across the country (Bonferroni corrected: *p <*0.005). These populations included Dodoma in central Tanzania; Tanga and Morogoro in the East; Kagera, Katavi, and Rukwa in the western part; and Ruvuma, Lindi, and Mtwara in southern Tanzania. However, where the overall HW tests were significant, the mean heterozygote deficits were generally mioderate (range: 0.407 - 0.660). Furthermore, HW tested for each locus separately within populations, showed that reduced *H*_S_ was not observed consistently across all loci (Additional file 4). Two populations, Pwani on the eastern coast and Kigoma in the west were found in Hardy-Weinberg equilibrium (Additional file 4).

### Population structure and differentiation

Fisher’s exact tests revealed highly significant genetic differentiation among all pairs of populations (*p*-values < 0.001 in all cases) (Table 1). As expected, the highest Chi-squared values were observed between the outgroup *An. parensis* population of Dodoma and *An. funestus* s.s. populations (range: 239.7 - 276.3). Additionally, among the *An. funestus* s.s. populations, Chi-square values of 276.3 were observed between the Kagera population in the north-west and the Lindi and Mtwara populations in southern Tanzania (Table 1). The lowest Chi-squared value of 101.4 was observed between Morogoro and Tanga, both of which located in the eastern part of Tanzania (Fig. 1).

**Table 1:** Pairwise estimates of genetic differentiation (using Fisher’s exact method) and genetic divergence (*F*_ST_) among *Anopheles fun*estus s.l. populations based on 12 microsatellite loci.

| Population | Dodoma | Pwani | Tanga | Kagera | Morogoro | Ruvuma | Kigoma | Katavi | Lindi | Mtwara |
| --- | --- | --- | --- | --- | --- | --- | --- | --- | --- | --- |
| **Pwani** | 261.1 | - |  |  |  |  |  |  |  |  |
| **Tanga** | 244.2 | 191.6 | - |  |  |  |  |  |  |  |
| **Kagera** | 250.5 | 218.0 | 205.0 | - |  |  |  |  |  |  |
| **Morogoro** | 252.1 | 190.5 | 101.4 | 189.8 | - |  |  |  |  |  |
| **Ruvuma** | 276.3 | 229.0 | 187.6 | 189.5 | 152.4 | - |  |  |  |  |
| **Kigoma** | 239.7 | 198.7 | 154.0 | 142.1 | 106.4 | 168.8 | - |  |  |  |
| **Katavi** | 265.4 | 248.4 | 238.6 | 155.5 | 165.3 | 186.6 | 138.4 | - |  |  |
| **Lindi** | 276.3 | 257.3 | 221.7 | 276.3 | 188.2 | 199.4 | 225.3 | 257.7 | - |  |
| **Mtwara** | 276.3 | 255.6 | 258.0 | 276.3 | 229.6 | 209.8 | 214.0 | 257.7 | 165.9 | - |
| **Rukwa** | 276.3 | 263.8 | 266.3 | 228.4 | 218.5 | 199.9 | 231.8 | 195.1 | 210.2 | 226.2 |
| **Pwani** | 0.2128 | - |  |  |  |  |  |  |  |  |
| **Tanga** | 0.2584 | 0.0717 | - |  |  |  |  |  |  |  |
| **Kagera** | 0.2414 | 0.1018 | 0.1549 | - |  |  |  |  |  |  |
| **Morogoro** | 0.2661 | 0.0735 | 0.0210 | 0.1385 | - |  |  |  |  |  |
| **Ruvuma** | 0.2961 | 0.1463 | 0.2005 | 0.0950 | 0.1662 | - |  |  |  |  |
| **Kigoma** | 0.2079 | 0.0596 | 0.0823 | 0.0418 | 0.0634 | 0.1045 | - |  |  |  |
| **Katavi** | 0.3074 | 0.0959 | 0.1663 | 0.0844 | 0.1293 | 0.1719 | 0.1108 | - |  |  |
| **Lindi** | 0.3777 | 0.1726 | 0.2279 | 0.2026 | 0.1683 | 0.2338 | 0.2036 | 0.1043 | - |  |
| **Mtwara** | 0.3268 | 0.1947 | 0.2881 | 0.2323 | 0.2587 | 0.2426 | 0.2089 | 0.2553 | 0.2655 | - |
| **Rukwa** | 0.3254 | 0.1291 | 0.2120 | 0.0852 | 0.1631 | 0.1662 | 0.1224 | 0.0361 | 0.0921 | 0.2637 |

\*The top matrix represents genetic differentiation based on Fisher’s exact method, while the bottom matrix represents $F_{ST}$ values. All Chi-squared and $F_{ST}$ values were associated with $p$ -values of less than 0.001. The Dodoma population represents *Anopheles parensis*.

### Genetic divergence and isolation by distance

Interspecific pairwise estimates of genetic divergence (*F*_ST_) between *An. parensis* from Dodoma and *An. funestus* s.s. populations averaged 0.282 (range: 0.208 - 0.378), with the highest between Dodoma and Lindi (0.378). Among *An. funestus* s.s. populations, the mean *F*_ST_ was 0.151 (range: 0.021 - 0.288) (Table 1). Moreover, geographically closer populations namely Morogoro, Tanga and Pwani in eastern Tanzania, and Kagera and Kigoma in western Tanzania had pairwise estimate values of *F*_ST_ below 0.1 (Fig. 1, Table 1). Interestingly, comparisons of genetic divergence among the 3 south-eastern populations (i.e., Lindi, Mtwara and Ruvuma) consistently yielded higher *F*_ST_ values indicating substantial genetic divergence despite their proximity (Fig. 1).

Considering *An. funestus* populations only, there was no correlation between linearised *F*_ST_ and log-transformed geographical distance (km) and thus no evidence of isolation-by-distance between populations (Mantel test: Adjusted *R*^2^ = -0.016, *p* = 0.569) (Fig. 3). Additionally, *F*_ST_ comparisons between pairs of populations located on either side of the Rift Valley were only marginally higher (mean = 0.18) than those involving pairs of populations on opposite sides of the rift (mean = 0.17) but this difference was not statistically significant (Kruskal-Wallis test: χ^2^ = 0.02, df = 1, *p* = 0.900).

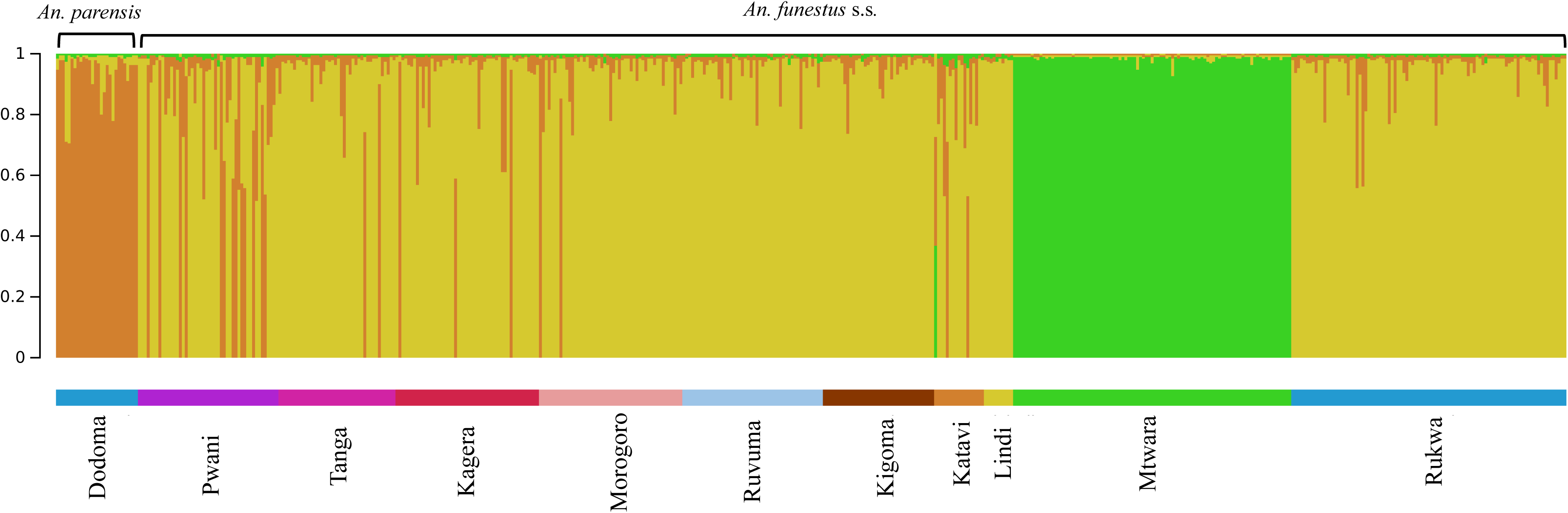

### Population clustering

The genetic relationships among populations were further analyzed using neighbor-joining cluster analysis, with *An. parensis* from Dodoma serving as an outgroup (Fig. 3). As anticipated, in the resulting tree topography, the *An. parensis* population from Dodoma was positioned at the most basal split in the resulting tree. Another subsequent major basal branch involved the *An. funestus* s.s. population from Mtwara, which separated distinctly from the other populations (Fig. 3).

The remaining nine *An. funestus* s.s. populations formed two primary clusters. The first cluster included populations from the western and southern parts of Tanzania, specifically Katavi, Rukwa, and Kagera in the west, and Lindi and Ruvuma in the south. The second cluster comprised populations from the eastern coast, including Tanga, Morogoro, and Pwani in the east, except for Kigoma located in the far west and thus geographically distant from those eastern populations (Fig. 1). Thus, the tree topology was not dominated by the effect of the Rift Valley, but rather by the lack of strong geographical clustering and by the distinctiveness of the southern Mtwara population.

### Gene flow and migration patterns

The genetic structure analysis using the Bayesian model-based clustering algorithm implemented in STRUCTURE, identified three distinct genetic clusters (K= 3) amongst our 11 populations. As expected, the *An. parensis* outgroup stood out from all *An. funestus* s.s. populations. Surprisingly, in all simulations, the Mtwara population consistently formed a distinct genetic cluster, clearly separated from the other populations and showing low levels of admixture with them (Fig. 4). This result corroborated the patterns of elevated genetic differentiation and divergence observed between Mtwara and other populations (Table 1), further highlighted by the neighbor-joining tree (Fig. 3).

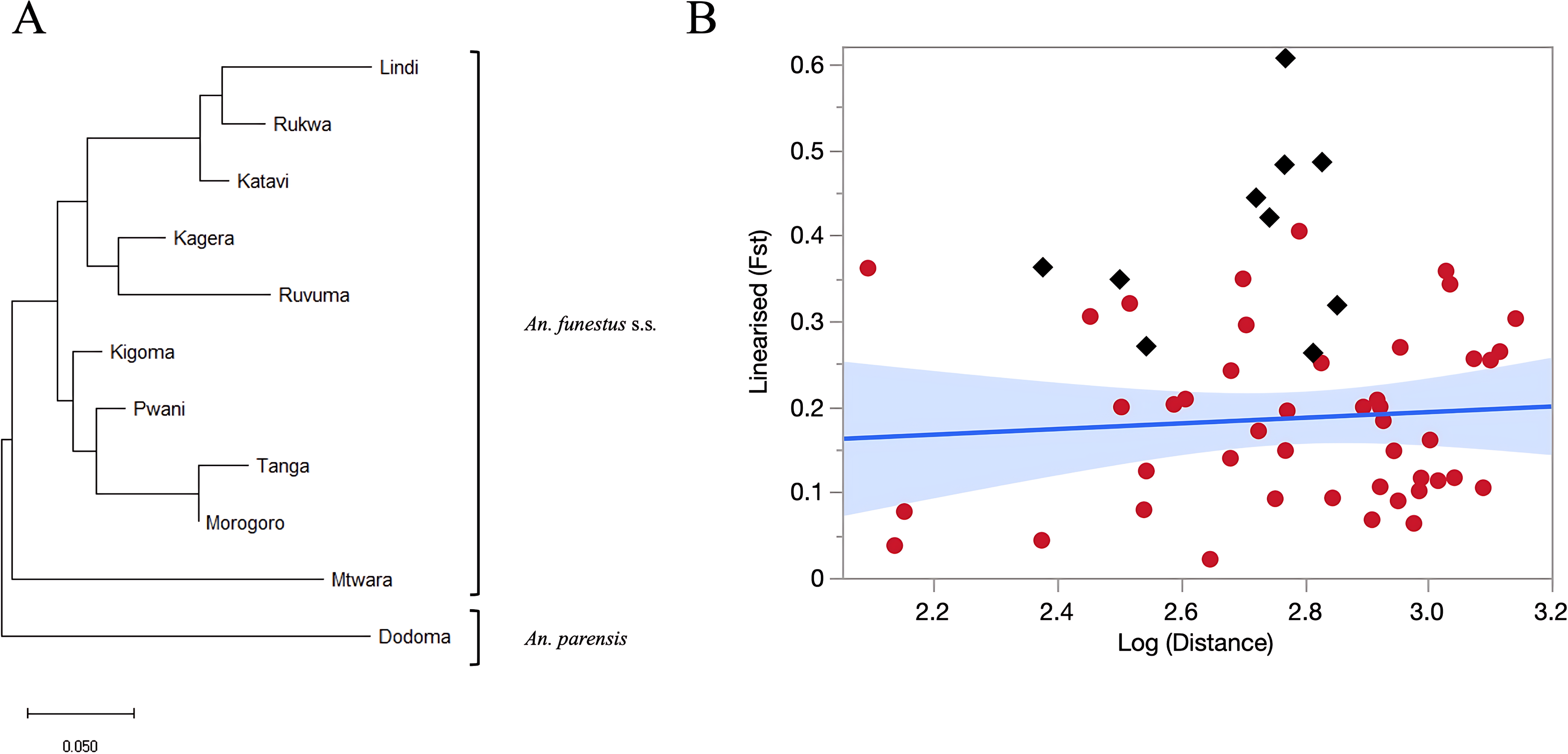

## Discussion

This study aimed at investigating population structure and gene flow within *An. funestus* s.s. populations across 10 administrative regions in mainland Tanzania. Here, for the first time, the same set of microsatellites previously used to study *An. funestus* populations across Africa [21] was successfully amplified in Tanzanian *An. funestus* s.s. populations, as well as in an *An. parensis* population used as an outgroup.

Pairwise comparisons of genetic differentiation and divergence confirmed that the *An. parensis* outgroup was more genetically distinct from the *An. funestus* s.s. populations than the latter were from each other. This was expected because of the distinct evolutionary pathways of these sibling species which are thought to have diverged from a common ancestor [40]. Geographically closer populations in eastern Tanzania, such as Morogoro, Tanga, and Pwani, and in western Tanzania, such as Kagera and Kigoma, exhibited the lowest pairwise *F*_ST_ values (less than 0.1), suggesting limited genetic divergence and high gene flow among these populations [41]. Interestingly, southeastern populations, including Lindi, Mtwara, and Ruvuma, showed higher pairwise *F*_ST_ values despite their geographical proximity.

Overall, the neighbour-joining clustering reflected these multifaceted patterns of isolation and gene flow. Aside from the *An. parensis* outgroup, the southern population of Mtwara also branched out basally from all other populations. This could be attributed to ecological, environmental, or anthropogenic factors that restrict gene flow and promote genetic divergence [41,42]. For instance, coastal southeastern Tanzania has a distinct ecological makeup, with Miombo woodlands, and mangrove ecosystems [43], which may limit mosquito dispersal. Additionally, soil properties such as texture, drainage, and organic content can influence the persistence of slow moving or permanent water bodies, which are preferred by *An. funestus* s.s. for larval development [44,45], These ecological constraints may further limit gene flow between regions. In that regard, Mahurunga village in Mtwara region, where sampling was conducted is located a few kilometers from the Indian Ocean coast on the estuary of the Rovuma River which defines the border with Mozambique. Thus, environmental conditions are strikingly different from those in the inland locations of Lindi and Ruvuma. Additionally, the Mtwara is the only collection site located right on the coast, and very close to the Mozambican border, and the R1260 road axis heading south into Mozambique. Given this location, it is plausible that the *An. funestus* population in Mtwara is genetically connected to other coastal populations, including those across the border in northern Mozambique. Coastal winds, human movement along the road corridor, and similar coastal habitats may facilitate passive or active dispersal. This connectivity could help explain the particularly high allelic diversity found in the Mtwara population compared to others.

The rest of the populations branched out into shallow clusters of the NJ tree: the first includes Tanga, Morogoro, Pwani, and Kigoma, while the second encompasses Lindi, Ruvuma, Katavi, Rukwa, and Kagera. Thus, the western population of Kigoma and the southern populations of Lindi and Ruvuma each clustered with populations that are geographically distant from them, rather than with their immediate neighbors. Although Kagera and Kigoma showed the lowest pairwise *F*_ST_ value, they appear in different clades in the *F*_ST_-based NJ tree. This pattern likely reflects their distinct relationships with other populations in the dataset, as the NJ algorithm clusters populations based on their genetic distances to all others, not just to each other. As a result, we did not find evidence for a strong effect of the Rift Valley on patterns of gene flow. The Rift valley that runs North-South across Tanzania is a major ecogeographical barrier which has been shown to reduce gene flow among *Anopheles* populations in east and southern Africa [46–48].

Unlike findings from other *Anopheles* species such as *An. gambiae* s.l. and *An. arabiensis*, where the Rift Valley often marks a clear genetic break, our results suggest that the Rift Valley does not strongly limit gene flow in *An. funestus* populations. This may reflect the species’ ecological requirements, tolerance for habitat variability, or possibly anthropogenic factors such as human-mediated dispersal along transportation routes like the central railway line. These factors may enable *An. funestus* to maintain gene flow across regions that are otherwise ecogeographically distinct. While there is no direct evidence that *An. funestus* is more frequently transported via anthropogenic means than other *Anopheles* species, its strong endophilic and anthropophilic tendencies may increase the likelihood of passive movement through human-associated environments, such as transport vehicles, cargo, or household materials. In regions where breeding sites are patchily distributed or isolated, such movement may help maintain gene flow between otherwise disconnected populations.

In this study, *F*_ST_ comparisons between populations on the same side of the Rift were only marginally lower than across the Rift. Furthermore, our NJ tree suggest extensive gene exchanges between some populations located on each side of the rift, possibly through a few major transportation axes. For example, Kigoma clustered with Pwani, two regions that have been historically connected by the central railway, the only East-West railway line in the country, which was completed in 1914 and runs to this day. This railway links Lake Tanganyika in the west to the central and eastern regions, facilitating freight and passenger transport, but possibly passive mosquito dispersal over long distances [49]. Variations in observed heterozygosity among populations further support this hypothesis, with comparatively higher levels of heterozygosity observed in the coastal region of Pwani and the western region of Kigoma than in all other populations.

Along the same lines, our collection sites from Lindi, Ruvuma and Rukwa were all located in the vicinity of a major East-West road axis used by trucks carrying goods from Tanzania to land-locked Zambia. This very busy road extends from the East Coast through large parts of the Lindi and Ruvuma in the Southeast, all the way to Zambian border and the nearby Southeastern region of Rukwa. Aside from the exceptions of Kigoma, Lindi and Ruvuma, most populations clustered together in a more predictable manner. Adjacent populations from eastern Tanzania, particularly along the coastal regions like Pwani and Tanga, grouped together suggesting high mosquito dispersal assisted by trade and migration along the eastern coast, combined with homogeneity in climate and ecology. Similarly, the clustering of the southern Ruvuma population with the western populations of Katavi and Rukwa may be explained by shared highland ecologies, combined important road connections. The distinctness of the Mtwara population, the overall moderate level or geographical genetic divergence among populations, combined with strong anthropogenic gene flow can explain the lack of isolation-by-distance observed among *An. funestus* s.s. populations in this study. These findings emphasize the importance of collection sites choice, particularly as it relates to road and rail axis, on population structure study outcomes. While microsatellite markers offer limited genomic resolution compared to genome wide SNPs, their high mutation rates make them particularly effective for detecting recent gene flow. This is especially relevant for understanding how resistance alleles may spread and how human movement along transport corridors influences mosquito populations. In our study, the observed clustering of distant populations such as Ruvuma and Rukwa, and the absence of isolation by distance, likely reflect recent gene flow facilitated by trade and human migration along major road networks. In contrast, a parallel whole genome SNP analysis using samples from the same populations [50] revealed more pronounced genomic divergence across geographic features such as the Rift Valley, suggesting patterns of local adaptation and historical separation over longer time scales. This comparison highlights the importance of marker choice, with microsatellites capturing more recent connectivity and SNPs revealing deeper evolutionary structure. Together, these approaches provide complementary insights into the demographic and evolutionary dynamics of *An. funestus* populations.

The possibility of population sub-structuring (cryptic taxa) within our populations was explored using classical Hardy-Weinberg tests, and Bayesian genotype assignment analyses with the software STRUCTURE. While statistically significant deviations from Hardy-Weinberg equilibrium (HWE) were observed in many populations these did not result from consistent deficits in heterozygotes (*Ho* < *Hs*) across loci. Therefore, they did not support sub-structuring and assortative mating. Instead, the random nature in the direction of these small deviations suggests that they were driven by excessive statistical power, due to large sample sizes (up to 96 mosquitoes per population) and relatively large number of polymorphic microsatellite loci. Next, the program STRUCTURE was used to identify possible sub-structuring within and between populations. This analysis confirmed the distinctiveness of the Mtwara population suggesting either a novel cryptic lineage, or an ecologically or geographically isolated populations. Strong reproductive isolation, leading to genetic divergence, is a common phenomenon in *Anopheles* species, where barriers like chromosomal forms [51], assortative mating [52], or ecological differences repeatedly resulted in speciation events in several well-known malaria vector complexes [51,53]. *An. funestus* is known for chromosomal inversions linked to adaptation and speciation, which may reduce recombination and promote reproductive isolation [54,55]. Further studies, including SNP genotyping or whole-genome sequencing along with testing of mating compatibility, will be needed to clarify if the distinctiveness of the Mtwara population truly reflects cryptic speciation or an unusually large intraspecific variation due to its distinct location. At present, the recently conducted sequencing NGS study that also included Mtwara, suggests that the later hypothesis is more likely than the former [50]. Overall genotype assignment mosaic plot generated by STRUCTURE was extremely clean with regard to distinctiveness of the Mtwara population, with no or very low level of admixture suggested with other populations. By comparison, the *An. parensis* outgroup was clearly outlined, but some genotypes from *An. funestus* s.s. populations suggested low levels of admixture between the two species, particularly in collections from the Pwani region. Thus some level of hybridization may occur between members of the Funestu*s* group, a possibility that remains to be formally tested, perhaps through sperm analyses of assortative mating as was conducted among members of the *An. gambiae* complex [56].

## Conclusions

These findings provide valuable insights into the genetic structure and gene flow of *An. funestus* s.s. populations in Tanzania. Despite regional variation in allele numbers and heterozygosity, genetic diversity was relatively uniform, and patterns of genetic divergence suggested high levels of population interconnectivity, even across the Rift Valley, likely due to passive mosquito dispersal along major road and railway axes. Notably, the Mtwara population displayed a distinct genetic profile, suggesting possible reproductive isolation, contrasted ecological pressures, and/or connectivity with Mozambican populations. Future research using NGS data should attempt to distinguish the effects of contemporary gene flow from those of local adaptation on population genetic structuring. This knowledge can inform the deployment of emerging vector control strategies, such as genetically modified mosquitoes, which may enhance future vector control efforts.

## Abbreviations

ITN: Insecticide treated bed net
IRS: Indoor residual spraying
PCR: Polymerase chain reaction
CDC: Centres for disease control and prevention
EDTA: Ethylene diamine tetra acetic acid
HWE: Hardy-Weinberg Equilibrium
Na: Number of alleles
Ra: Allelic richness

## Acknowledgments

We extend our heartfelt gratitude to the community members across all districts for their participation in this study. We also sincerely thank Mr. Simon Ashall, Dr. Florian Noulin at Keele University, and Mr. Alex Limwagu, Mr. Augusto Mwambaluka, and Mr. Jonael Msangi at the Ifakara Health Institute for their invaluable support and contributions. We are especially grateful to Prof. Igor Sharakhov for kindly providing the original polytene chromosome image, which was used, with permission, as Figure 2 in this manuscript.

## Availability of data and materials

All data supporting the conclusions of this article are provided within the text.

## Authors’ contributions

IHN, GM, HB, JOO, JPM, EWK, NJG, NFK, and EEH contributed to the planning and implementation of field activities, including coordinating logistics, supervising sample collection teams, and ensuring adherence to ethical and sampling protocols. SAM, PAK and BS performed all laboratory analyses. SAM, NJG, RG, FOO, BS, and FT contributed to the study design and interpretation of results. SAM, BS, and FT drafted the manuscript with input from FOO. All authors read, revised, and approved the final manuscript.

## Ethics approval and consent to participate

Ethical approvals for this project were obtained from Ifakara Health Institute’s Institutional Review Board (Protocol ID: IHI/IRB/No: 19 – 2017 and IHI/IRB/EXT/No: 33 - 2022) and the Medical Research Coordinating Committee (MRCC) at the National Institute for Medical Research, in Tanzania (Protocol ID: NIMR/HQ/R.8a/Vol.IX/2697 and NIMR/HQ/R.8a/Vol.IX/3494). Written consents were sought from all participants of this study, after they had understood the purpose and procedure of the discussions.

## Consent for publication

Permission to publish this study was obtained from National Institute for Medical Research, in Tanzania (BD.242/437/01C/135).

