## Additional file 1 for "Microsatellite Analysis of Genetic Differentiation Among Populations of the Malaria Vector, *Anopheles funestus*, across Mainland Tanzania Reveals Contrasted Patterns of Geographic Isolation and Gene Flow"

| Region | Village | Population | Geographical coordinates | Altitude | Great Rift Valley |
| --- | --- | --- | --- | --- | --- |
| Kagera | Byeju | *An. funestus* s.s | 1.07892°S, 31.36819°E | 1196 | Western |
| Kigoma | Kitahana | *An. funestus* s.s | 3.62452°S, 30.68260°E | 874 | Western |
| Kigoma | Kakonko | *An. funestus* s.s | 3.2799°S, 30.9661°E | 1260 | Western |
| Katavi | Vikonge | *An. funestus* s.s | 6.08851°S, 30.93067°E | 1800 | Western |
| Rukwa | Kabwe | *An. funestus* s.s | 7.0437°S, 30.5553°E | 1182 | Western |
| Dodoma | Dabalo | *An. parensis* | 6.1811°S, 35.7469°E | 1124 | Eastern |
| Tanga | Kwamwale | *An. funestus* s.s | 5.16844°S, 38.72226°E | 200 | Eastern |
| Pwani | Mwavi | *An. funestus* s.s | 6.40882°S, 38.58196°E | 10 | Eastern |
| Morogoro | Igumbiro | *An. funestus* s.s | 8.35021°S, 36.67299°E | 340 | Eastern |
| Morogoro | Sululu | *An. funestus* s.s | 8.00324°S, 36.83118°E | 315 | Eastern |
| Lindi | Mtua Longa | *An. funestus* s.s | 10.21532°S, 39.49712°E | 300 | Eastern |
| Mtwara | Mahurunga | *An. funestus* s.s | 10.32186°S, 40.16386°E | 900 | Eastern |
| Ruvuma | Nandembo | *An. funestus* s.s | 10.95887°S, 37.23666°E | 1200 | Eastern |

Additional file 1: Sampling sites of *Anopheles funestus* mosquitoes in Tanzania with geographical attributes
