## Additional file 2 for "Microsatellite Analysis of Genetic Differentiation Among Populations of the Malaria Vector, *Anopheles funestus*, across Mainland Tanzania Reveals Contrasted Patterns of Geographic Isolation and Gene Flow"

**Additional file 2:** Eighteen microsatellite markers previously used in population genetic studies of *Anopheles funestus* mosquitoes and their multiplex design

| Triplexes | Locus | Accession Number | Repeat Motif | Sequence (5′ - 3′) | Size range |
| --- | --- | --- | --- | --- | --- |
| 1 | AFND12 | *AF171042* | (GA)_6_ | ATTO565- GTTCTCCATCGCTGTTCTACTC  R : TATAACGTTTCGTACACACGCC | 87-107 |
|  | AFUB6 | *AY029713* | (CTG)2+2+4+4+3 | YAKYE- CCAGCAGGTGTGGAGGAC | 145-151 |
|  |  |  |  | R : GTCGTACAAAAGCACCACCA |  |
|  | AFND6 | *AF171036* | (GT)7+3 | FAM- GCTTCTTCTCCCCTAATCTG | 184-212 |
|  |  |  |  | R : TCCTGCTTTTTAGTTTGTCG |  |
| 2 | AFND7 | *AF171037* | (GT)_8_ | ATTO565- TGCATCATTCGACTCGGAAG | 70-84 |
|  |  |  |  | R: AACGGCACTACCGTTCACTG |  |
|  | AFND32 | *AY291371* | (TG)_2_TT(TG)_3_AG(TG)_6_ | YAKYE- GAAGCATTTTGGGTTAGACTC | 103-121 |
|  |  |  |  | R: GCAGTTGTTTACCTTTCACTG |  |
|  | AFND5 | *AF171035* | (GT)5+3 | FAM- CCTCTCGTTGTGTTGCCTAC | 169-183 |
|  |  |  |  | R : GTTCATACGTTGCCCGATTT |  |
| 3 | AFUB12 | *AY029719* | (AGG)_7_(TG)_4_ | ATTO565-TGGGGAACTGGTCGTTAGAG | 87-107 |
|  |  |  |  | R: CTGGTGATGGGATTGAGGAT |  |
|  | Fun R | *AY6022* | (TG)_8_ T(TG)_3_ | YAKYE- GTAGTCGATGGTGCCGTGTG | 132-148 |
|  |  |  |  | R: ACCGTCCCTTCCATCTGTGA |  |
|  | AFUB3 | *AY029710* | (CAG)_2+3+2_ | FAM- GGGAAGGATTCGACCTTAGC | 164-182 |
|  |  |  |  | R: GCCGCCATTTAGTAGCAGTT |  |
| 4 | AFND30 | *AY291369* | (TG)_9_CC(TG)_2_ | ATTO565- GTTCAGCTGTTGGTGTGTTAG  R: TTTTCGTACGGAGAAAAATG | 81-107 |
|  | Fun O | *AY6019* | (CA)_6_ TA(AC)_4_ | YAKYE- GCACACATTTCAGGCAGC  R: GCCCACATTCTGCACCTT | 116-124 |
|  | AFND19 | *AF171049* | (GT)12 | FAM- CAGAACCACTTCGATTCAAC  R: CCTGCACTCAGAAACACAC | 172-205 |
| 5 | Fun F | *AY6010* | (TG)_9_ | ATTO565- GCCTTCAGTTTCGATTGGCG | 104-118 |
|  |  |  |  | R: AATAAGATGCGACCGTGGC |  |
|  | AFUB11 | *AY029718* | (CTG)_3+5+2+2_ | YAKYE- CAGTTTCTGCGTGGAGGAAT | 188-195 |
|  |  |  |  | R: AGCAGCTGATGAGCCATCTC |  |
|  | AFUB10 | *AY029717* | (GCT)_3+2+4+5_ | FAM- TGTCCATGTACAACCGCAAC | 195-210 |
|  |  |  |  | R: TTCTCCAGCATCATCAGCAC |  |
| 6 | Fun L | *AY6016* | (GT) _8_ | ATTO565- AACAGTGGAAGGCAAATTGC | 140-166 |
|  |  |  |  | R: GCACGGTTACCACTGCTCA |  |
|  | AFND40 | *AY291374* | (AC)_7_..(17nts)..(AC)_6_ | YAKYE- ATTCATCCTGTGATGCTTTG | 188-200 |
|  |  |  |  | R: AGGCTCTTCTTTGCACTGT |  |
|  | Fun Q | *AY6021* | (TG) _9_ | FAM- GCAAACTGCTAGTAAATGTTTCC | 238-246 |
|  |  |  |  | R: ACATTTCCACAATTTGCGC |  |
