## Additional file 3 for "Microsatellite Analysis of Genetic Differentiation Among Populations of the Malaria Vector, *Anopheles funestus*, across Mainland Tanzania Reveals Contrasted Patterns of Geographic Isolation and Gene Flow"

**Additional file 3:** Hardy-Weinberg equilibrium results of 18 microsatellite markers in *Anopheles parensis* (Dodoma) and *Anopheles funestus* s.s. mosquitoes from Tanzania Mainland

| Locus | Chromosomal location |  | Dodoma  (N=28) | Pwani  (N=48) | Tanga  (N=40) | Kagera  (N=48) | Morogoro  (N=48) | Ruvuma  (N=48) | Kigoma  (N=38) | Katavi  (N=96) | Lindi  (N=96) | Mtwara  (N=96) | Rukwa  (N=96) |
| --- | --- | --- | --- | --- | --- | --- | --- | --- | --- | --- | --- | --- | --- |
| AFND12 | X | Na | 7 | 2 | 2 | 6 | 2 | 2 | 6 | 6 | 1 | 2 | 2 |
|  |  | HW | 0.018 | 0.229 | 0.162 | 0.012 | - | 1.000 | 0.008 | - | - | - | 1.000 |
| FUN Q | X | Na | 3 | 5 | 5 | 7 | 5 | 7 | 9 | 6 | 4 | 18 | 6 |
|  |  | HW | 0.100 | **0.000*** | **0.000*** | 0.005 | **0.000*** | 0.106 | 1.000 | **0.000*** | 0.391 | **0.000*** | 1.000 |
| AFUB3 | 2R (inversion) | Na | 11 | 5 | 9 | 7 | 4 | 7 | 11 | 8 | 7 | 8 | 8 |
|  |  | HW | **0.000*** | 0.191 | 0.646 | 0.674 | 0.495 | 0.916 | 0.079 | 0.073 | 0.783 | 0.148 | 0.083 |
| AFND40 | 2R (inversion) | Na | 7 | 4 | 5 | 6 | 4 | 3 | 6 | 5 | 3 | 13 | 4 |
|  |  | HW | 0.003 | **0.001*** | 0.132 | 0.033 | **0.000*** | 0.531 | 0.988 | **0.000*** | **0.000*** | **0.000*** | 0.010 |
| AFUB6 | 2R (inversion) | Na | 7 | 3 | 3 | 3 | 6 | 3 | 5 | 3 | 4 | 2 | 3 |
|  |  | HW | **0.000*** | 0.046 | 0.526 | 0.011 | 0.004 | 0.054 | 0.125 | **0.000*** | **0.001*** | **0.000*** | 1.000 |
| FUN R | 2R (inversion) | Na | 7 | 6 | 3 | 4 | 3 | 2 | 7 | 6 | 4 | 4 | 6 |
|  |  | HW | **0.000*** | **0.001*** | 0.007 | **0.000*** | 1.000 | **0.000*** | 0.872 | **0.000*** | 1.000 | 0.358 | 1.000 |
| AFND6 | 2R (inversion) | Na | 10 | 7 | 11 | 9 | 6 | 10 | 11 | 7 | 4 | 8 | 10 |
|  |  | HW | 0.030 | 0.064 | **0.000*** | 0.030 | 0.038 | 0.113 | **0.002*** | 0.091 | 0.009 | **0.000*** | 0.599 |
| AFND30 | 2R (inversion) | Na | 9 | 8 | 7 | 12 | 6 | 5 | 11 | 11 | 5 | 12 | 11 |
|  |  | HW | 0.081 | 0.075 | **0.000*** | 0.672 | **0.000*** | 0.086 | **0.000*** | **0.000*** | 0.409 | **0.000*** | 0.032 |
| AFND5 | 2R (inversion) | Na | 6 | 5 | 8 | 4 | 5 | 6 | 8 | 3 | 10 | 3 | 7 |
|  |  | HW | 0.010 | 0.486 | **0.001*** | 0.064 | **0.000*** | 0.820 | 0.255 | 1.000 | **0.000*** | 0.009 | 0.908 |
| AFND32 | 2R (inversion) | Na | 8 | 8 | 10 | 9 | 9 | 6 | 8 | 10 | 6 | 8 | 7 |
|  |  | HW | 0.031 | 0.311 | **0.001*** | 0.085 | 0.171 | 0.435 | 0.847 | 0.019 | **0.000*** | **0.000*** | 0.024 |
| FUN O | 2R | Na | 6 | 7 | 5 | 9 | 6 | 5 | 9 | 7 | 4 | 7 | 6 |
|  |  | HW | **0.001*** | 0.130 | 0.276 | 0.118 | 0.520 | 0.193 | 0.515 | **0.000*** | 0.116 | 0.019 | 0.645 |
| AFUB10 | 2L | Na | 8 | 10 | 6 | 8 | 8 | 6 | 9 | 6 | 7 | 9 | 7 |
|  |  | HW | **0.000*** | 0.079 | 0.208 | **0.001*** | 0.534 | 0.006 | 0.381 | **0.000*** | 0.025 | 0.749 | 0.870 |
| FUN L | 2L | Na | 9 | 9 | 11 | 12 | 10 | 8 | 17 | 14 | 7 | 4 | 13 |
|  |  | HW | **0.001*** | 0.726 | 0.007 | 0.352 | 0.678 | 0.540 | 0.920 | **0.000*** | 0.068 | **0.000*** | 0.060 |
| AFUB11 | 2L | Na | 3 | 3 | 5 | 4 | 5 | 8 | 5 | 3 | 5 | 7 | 7 |
|  |  | HW | 0.050 | **0.002*** | 0.004 | **0.000*** | **0.002*** | **0.000*** | 0.434 | **0.001*** | 0.013 | **0.002*** | **0.000*** |
| AFND7 | 3R | Na | 6 | 8 | 8 | 7 | 7 | 7 | 13 | 8 | 3 | 6 | 6 |
|  |  | HW | 0.839 | 0.652 | 0.036 | **0.000*** | 0.142 | 0.497 | 0.541 | **0.000*** | **0.000*** | 0.009 | 0.175 |
| AFND19 | 3R (inversion) | Na | 15 | 11 | 6 | 8 | 8 | 9 | 9 | 10 | 7 | 11 | 9 |
|  |  | HW | **0.000*** | 0.210 | 0.061 | 0.072 | 0.071 | 0.609 | 0.410 | **0.000*** | **0.000*** | **0.000*** | 0.180 |
| AFUB12 | 3L | Na | 8 | 9 | 7 | 2 | 3 | 2 | 2 | 4 | 3 | 5 | 3 |
|  |  | HW | 0.012 | **0.000*** | 0.035 | 0.005 | **0.001*** | 0.527 | **0.000*** | **0.000*** | 0.004 | **0.000*** | 0.216 |
| FUN F | 3L (inversion) | Na | 7 | 7 | 7 | 7 | 6 | 9 | 6 | 6 | 3 | 6 | 9 |
|  |  | HW | **0.001*** | 1.000 | 0.395 | 0.228 | 0.090 | 0.232 | 0.284 | 0.004 | **0.000*** | 0.041 | 0.415 |
| Mean across all loci | | Na | 7.6 | 6.5 | 6.6 | 6.9 | 5.7 | 5.8 | 8.4 | 6.8 | 4.8 | 7.4 | 6.9 |

*Bold p-values with asterisk are statistically significant by Bonferroni correction (*p* < 0.05/18 = 0.0028).
