## Additional file 4 for "Microsatellite Analysis of Genetic Differentiation Among Populations of the Malaria Vector, *Anopheles funestus*, across Mainland Tanzania Reveals Contrasted Patterns of Geographic Isolation and Gene Flow"

**Additional file 4:** Genetic variability at 12 microsatellite loci in *Anopheles parensis* (Dodoma) and *Anopheles funestus* s.s. from Tanzania mainland

| *An. parensis* | | | | | *Anopheles funestus* s.s. | | | | | | | | | |  |
| --- | --- | --- | --- | --- | --- | --- | --- | --- | --- | --- | --- | --- | --- | --- | --- |
| Locus | Chromosomal  location | |  | Dodoma  N = 28 | Pwani  N = 48 | Tanga  N = 40 | Kagera  N = 48 | Morogoro  N = 48 | Ruvuma  N = 48 | Kigoma  N = 38 | Katavi  N = 96 | Lindi  N = 96 | Mtwara  N = 96 | Rukwa  N = 96 | Mean^Af^  N = 682 |
| AFND12 | X | | Na | 7 | 2 | 2 | 6 | 2 | 2 | 6 | 6 | 1 | 2 | 2 | 0.493 |
|  |  |  | *H*o | 0.240 | 0.604 | 0.500 | 0.375 | 0.000 | 0.021 | 0.211 | 0.104 | 0.000 | 0.010 | 0.011 | 0.184 |
|  |  |  | *H*_S_ | 0.228 | 0.440 | 0.403 | 0.411 | 0.045 | 0.021 | 0.200 | 0.149 | 0.000 | 0.010 | 0.011 | 0.169 |
|  |  |  | *F*_IS_ | -0.055 | -0.373 | -0.242 | 0.087 | 1 | 0.000 | -0.055 | 0.301 | NA | 0.000 | 0.000 | 0.229 |
| FUN Q | X | | Na | 3 | 5 | 5 | 7 | 5 | 7 | 9 | 6 | 4 | 18 | 6 | 7.200 |
|  |  |  | *H*o | 0.357 | 0.875 | 0.875 | 0.458 | 0.563 | 0.854 | 0.711 | 0.196 | 0.183 | 0.404 | 0.042 | 0.516 |
|  |  |  | *H*_S_ | 0.308 | 0.708 | 0.553 | 0.695 | 0.591 | 0.639 | 0.593 | 0.185 | 0.180 | 0.543 | 0.246 | 0.493 |
|  |  |  | *F*_IS_ | -0.159 | -0.235 | -0.583 | 0.340 | 0.048 | -0.338 | -0.198 | -0.060 | -0.017 | 0.256 | 0.829 | 0.290 |
| AFND40 | 2R (inversion) | | Na | 7 | 4 | 5 | 6 | 4 | 3 | 6 | 5 | 3 | 13 | 4 | 5.300 |
|  |  |  | *H*o | 0.607 | 0.438 | 0.850 | 0.638 | 0.729 | 0.625 | 0.763 | 0.646 | 0.366 | 0.968 | 0.958 | 0.698 |
|  |  |  | *H*_S_ | 0.798 | 0.675 | 0.647 | 0.749 | 0.655 | 0.656 | 0.714 | 0.642 | 0.589 | 0.677 | 0.545 | 0.655 |
|  |  |  | *F*_IS_ | 0.239 | 0.352 | -0.314 | 0.148 | -0.113 | 0.047 | -0.069 | -0.006 | 0.379 | -0.431 | 0.239 | 0.210 |
| FUN R | 2R (inversion) | | Na | 7 | 6 | 3 | 4 | 3 | 2 | 7 | 6 | 4 | 4 | 6 | 4.500 |
|  |  |  | *H*o | 0.667 | 0.818 | 0.075 | 0.298 | 0.042 | 0.083 | 0.447 | 0.526 | 0.115 | 0.531 | 0.387 | 0.332 |
|  |  |  | *H*_S_ | 0.541 | 0.740 | 0.167 | 0.475 | 0.157 | 0.081 | 0.737 | 0.536 | 0.110 | 0.464 | 0.672 | 0.414 |
|  |  |  | *F*_IS_ | -0.232 | -0.106 | 0.550 | 0.373 | 0.735 | -0.033 | 0.393 | 0.018 | -0.037 | -0.146 | 0.424 | 0.282 |
| AFND30 | 2R (inversion) | | Na | 9 | 8 | 7 | 12 | 6 | 5 | 11 | 11 | 5 | 12 | 11 | 8.800 |
|  |  |  | *H*o | 0.679 | 0.870 | 0.650 | 0.625 | 0.792 | 0.404 | 0.684 | 0.447 | 0.522 | 0.563 | 0.417 | 0.597 |
|  |  |  | *H*_S_ | 0.859 | 0.801 | 0.691 | 0.865 | 0.720 | 0.700 | 0.838 | 0.873 | 0.417 | 0.750 | 0.791 | 0.745 |
|  |  |  | *F*_IS_ | 0.210 | -0.085 | 0.059 | 0.277 | -0.099 | 0.423 | 0.183 | 0.488 | -0.252 | 0.250 | 0.473 | 0.259 |
| AFND32 | 2R (inversion) | | Na | 8 | 8 | 10 | 9 | 9 | 6 | 8 | 10 | 6 | 8 | 7 | 8.100 |
|  |  |  | *H*o | 0.571 | 0.696 | 0.725 | 0.583 | 0.708 | 0.674 | 0.763 | 0.729 | 0.484 | 0.563 | 0.762 | 0.669 |
|  |  |  | *H*_S_ | 0.739 | 0.773 | 0.799 | 0.804 | 0.790 | 0.726 | 0.766 | 0.771 | 0.671 | 0.703 | 0.681 | 0.748 |
|  |  |  | *F*_IS_ | 0.227 | 0.100 | 0.092 | 0.275 | 0.104 | 0.071 | 0.004 | 0.054 | 0.280 | 0.200 | -0.118 | 0.130 |
| FUN O | 2R | | Na | 6 | 7 | 5 | 9 | 6 | 5 | 9 | 7 | 4 | 7 | 6 | 6.500 |
|  |  |  | *H*o | 0.786 | 0.896 | 0.625 | 0.563 | 0.563 | 0.457 | 0.737 | 0.583 | 0.646 | 0.448 | 0.400 | 0.592 |
|  |  |  | *H*_S_ | 0.665 | 0.736 | 0.616 | 0.583 | 0.642 | 0.547 | 0.746 | 0.636 | 0.556 | 0.576 | 0.554 | 0.619 |
|  |  |  | *F*_IS_ | -0.181 | -0.218 | -0.015 | 0.036 | 0.124 | 0.165 | 0.012 | 0.082 | -0.161 | 0.222 | 0.278 | 0.131 |
| AFUB10 | 2L | | Na | 8 | 10 | 6 | 8 | 8 | 6 | 9 | 6 | 7 | 9 | 7 | 7.600 |
|  |  |  | *H*o | 0.778 | 0.542 | 0.475 | 0.773 | 0.468 | 0.646 | 0.889 | 0.617 | 0.457 | 0.708 | 0.370 | 0.595 |
|  |  |  | *H*_S_ | 0.845 | 0.767 | 0.593 | 0.826 | 0.709 | 0.621 | 0.784 | 0.697 | 0.561 | 0.641 | 0.665 | 0.686 |
|  |  |  | *F*_IS_ | 0.079 | 0.293 | 0.198 | 0.064 | 0.340 | -0.041 | -0.134 | 0.115 | 0.187 | -0.105 | 0.443 | 0.192 |
| FUN L | 2L | | Na | 9 | 9 | 11 | 12 | 10 | 8 | 17 | 14 | 7 | 4 | 13 | 10.500 |
|  |  |  | *H*o | 0.963 | 0.857 | 0.795 | 0.681 | 0.809 | 0.702 | 1.000 | 0.787 | 0.596 | 1.000 | 0.337 | 0.756 |
|  |  |  | *H*_S_ | 0.769 | 0.871 | 0.740 | 0.627 | 0.686 | 0.724 | 0.857 | 0.786 | 0.684 | 0.589 | 0.587 | 0.715 |
|  |  |  | *F*_IS_ | -0.255 | 0.005 | -0.081 | -0.097 | -0.154 | 0.022 | -0.166 | -0.007 | 0.138 | -0.698 | 0.414 | 0.178 |
| AFUB11 | 2L | | Na | 3 | 3 | 5 | 4 | 5 | 8 | 5 | 3 | 5 | 7 | 7 | 5.200 |
|  |  |  | *H*o | 0.217 | 0.413 | 0.525 | 0.488 | 0.354 | 0.400 | 0.421 | 0.511 | 0.564 | 0.531 | 0.473 | 0.468 |
|  |  |  | *H*_S_ | 0.621 | 0.582 | 0.625 | 0.480 | 0.616 | 0.540 | 0.640 | 0.423 | 0.655 | 0.645 | 0.583 | 0.579 |
|  |  |  | *F*_IS_ | 0.650 | 0.290 | 0.160 | -0.018 | 0.425 | 0.259 | 0.343 | -0.209 | 0.139 | 0.177 | 0.189 | 0.221 |
| AFND19 | 3R (inversion) | | Na | 15 | 11 | 6 | 8 | 8 | 9 | 9 | 10 | 7 | 11 | 9 | 8.800 |
|  |  |  | *H*o | 0.815 | 0.875 | 0.400 | 0.596 | 0.688 | 0.625 | 0.842 | 0.792 | 0.547 | 0.800 | 0.698 | 0.686 |
|  |  |  | *H*_S_ | 0.875 | 0.739 | 0.474 | 0.704 | 0.718 | 0.633 | 0.714 | 0.760 | 0.705 | 0.736 | 0.738 | 0.692 |
|  |  |  | *F*_IS_ | 0.068 | -0.183 | 0.156 | 0.154 | 0.042 | 0.013 | -0.179 | -0.042 | 0.223 | -0.087 | 0.054 | 0.113 |
| AFUB12 | | 3L | Na | 8 | 9 | 7 | 2 | 3 | 2 | 2 | 4 | 3 | 5 | 3 | 4.000 |
|  |  |  | *H*o | 0.429 | 0.596 | 0.250 | 0.333 | 0.302 | 0.229 | 0.447 | 0.323 | 0.404 | 0.280 | 0.183 | 0.335 |
|  |  |  | *H*_S_ | 0.434 | 0.609 | 0.571 | 0.508 | 0.514 | 0.458 | 0.507 | 0.521 | 0.472 | 0.528 | 0.408 | 0.510 |
|  |  |  | *F*_IS_ | 0.012 | 0.021 | 0.562 | 0.343 | 0.412 | 0.500 | 0.118 | 0.38 | 0.144 | 0.470 | 0.552 | 0.350 |
| Mean across all loci | | | Na | 7.500 | 6.833 | 6.000 | 7.250 | 5.750 | 5.250 | 8.167 | 7.333 | 4.667 | 8.333 | 6.750 | 6.416 |
|  |  |  | *H*o | 0.592 | 0.707 | 0.562 | 0.534 | 0.502 | 0.477 | 0.660 | 0.522 | 0.407 | 0.567 | 0.420 | 0.536 |
|  |  |  | *H*_S_ | 0.640 | 0.703 | 0.573 | 0.644 | 0.570 | 0.529 | 0.675 | 0.582 | 0.467 | 0.572 | 0.540 | 0.585 |
|  |  |  | *F*_IS_ | 0.197 | 0.188 | 0.251 | 0.184 | 0.300 | 0.159 | 0.155 | 0.147 | 0.178 | 0.254 | 0.334 | 0.215 |
| Overall HWE | | | | **0.000^*^** | 0.053 | **0.000^*^** | **0.000^*^** | **0.000^*^** | **0.002^*^** | 0.020 | **0.000^*^** | **0.000^*^** | **0.000^*^** | **0.000^*^** |  |

*P-values in bold and with asterisk are statistically significant after Bonferroni correction (*p* < 0.05/11 = 0.005). Mean^Af^ represents overall mean values per locus across all *An. funestus* s.s. populations.
